# *Campylobacter jejuni* transcriptome changes during loss of culturability in water

**DOI:** 10.1101/150383

**Authors:** Christina Bronowski, Kasem Mustafa, Ian B. Goodhead, Chloe E. James, Charlotte Nelson, Anita Lucaci, Paul Wigley, Tom J. Humphrey, Nicola J. Williams, Craig Winstanley

## Abstract

**Background:** The natural environment serves as a potential reservoir for *Campylobacter,* the leading cause of bacterial gastroenteritis in humans. However, little is understood about the mechanisms underlying variations in survival characteristics between different strains of *C. jejuni* in natural environments, including water.

**Results:** We identified three *Campylobacter jejuni* strains that exhibited variability in their ability to retain culturability after suspension in water at two different temperatures (4°C and 25°C). Of the three, strains *C. jejuni* M1 exhibited the most rapid loss of culturability whilst retaining viability. Using RNAseq transcriptomics, we characterised C. *jejuni* M1 gene expression in response to suspension in water by analyzing bacterial suspensions recovered immediately after introduction into water (Time 0), and from two sampling time/temperature combinations where considerable loss of culturability was evident, namely (i) after 24 h at 25°C, and (ii) after 72 h at 4°C. Transcript data were compared with a culture-grown control. Some gene expression characteristics were shared amongst the three populations recovered from water, with more genes being up-regulated than down. Many of the up-regulated genes were identified in the Time 0 sample, whereas the majority of down-regulated genes occurred in the 25°C (24 h) sample.

**Conclusions:** Variations in expression were found amongst genes associated with oxygen tolerance, starvation and osmotic stress. However, we also found upregulation of flagellar assembly genes, accompanied by down-regulation of genes involved in chemotaxis. Our data also suggested a switch from secretion via the *sec* system to via the *tat* system, and that the quorum sensing gene *luxS* may be implicated in the survival of strain M1 in water. Variations in gene expression also occurred in accessory genome regions. Our data suggest that despite the loss of culturability, *C. jejuni* M1 remains viable and adapts via specific changes in gene expression.

## Background

*Campylobacter jejuni* is the most common bacterial cause of gastroenteritis in Europe and the USA [1-3]. Although campylobacteriosis is primarily considered to be a zoonotic infection, mostly transmitted to humans through consumption of contaminated poultry [4], it is clear that the environment can play a role in transmission either directly, for example via unchlorinated drinking water [5], or indirectly, via farm animals that acquire the pathogen from the environment [4, 6]. Indeed, contaminated groundwater is considered to be a potential source of transmission of *Campylobacter* to poultry flocks and to humans directly [7-15].

The survival mechanisms employed by *Campylobacter* outside of the host are not well understood. Genomic studies suggest that the ability of *C. jejuni* to regulate gene expression in response to environmental stresses may be limited because of the lack of many of the stress response mechanisms possessed by other bacterial species [6, 16, 17]. However, despite its apparent limited ability to respond to stress, *Campylobacter* can survive in the environment and remain infectious [18]. It has been suggested that *Campylobacter spp.* can survive in natural water by entering a viable but non culturable (VBNC) state [19]. During the VBNC state, *Campylobacter* form coccoid-shaped cells with an intact cell membrane, which remain viable according to various measures of metabolic activity, but are unable to grow on routine culture media [20]. Bacteria in this state are capable of causing infections or colonising a host and can be returned to a state of culturability [21].

Previous studies have shown that survival of *Campylobacter* in natural water samples or ground water is highly dependent upon temperature and strain origin [22-24]. Low temperatures (around 4°C) enhance the survival of *Campylobacter,* whereas at increased temperatures (20-25°C) viability declines rapidly [22, 25]. Survival times for different C. *jejuni* isolates in water can vary from a few days at ambient temperatures to four months at 4°C [26].

Many factors could contribute to the variations observed between *Campylobacter* strains with respect to survival in water [18]. These include the ability to metabolize available nutrients and to deal with oxidative and osmotic stress [25, 27]. Previous epidemiological studies have suggested that some strain types, defined using schemes such as Multi Locus Sequence Typing (MLST), are more commonly found in the environment [28-30]

There are significant gaps in our knowledge concerning variations in the survival of different *C. jejuni* strain types in the environment. These could be due to differences either in gene content or in gene expression. Variations in the gene content of *C. jejuni* strains, linked to virulence potential, have been described previously [31]. These include variations in surface structures, such as glycosylation patterns of flagellin or the structures of lipooligosaccharides, but also include metabolic traits [32]. Genes supporting oxygen-independent respiration and the catabolism of amino acids and peptides are particularly over-represented in strains that are robust colonisers of poultry, compared to strains that do not colonise well [30, 31, 33-36]. Little is known about how variations in gene expression contribute to the ability to survive in the environment. A better understanding of the occurrence and behaviour of *Campylobacter* in water, and the mechanisms underlying variations has implications for food safety and public health [37].

In this study we demonstrate how strains of *C. jejuni* adjust differently to an aquatic lifestyle by comparing the culturability of *C. jejuni* strains after prolonged exposure to water. We show that *C. jejuni* strain M1, though rapidly losing culturability on standard laboratory media, retains viability in water and, using strand-specific Illumina RNA Seq analysis, we identify gene expression changes occurring during the survival process.

## Results and Discussion

### Loss of culturability during survival in water

In this study, we focused on three *C. jejuni* strains exhibiting differences in their ability to retain culturability during incubation in water at specific time points and temperatures (24 h at 25°C and 72 h at 4°C). The ability to retain culturability in water was tested using three biological replicates for each of the three strains of *C. jejuni:* M1 (ST137, clonal complex ST45, associated with severe human infection) [38], 1336 (ST841, a representative of the water/wild-life clade)[30], and strain 414 (ST3704, associated with bank voles) [39]. Clear variations were observed between the three strains with respect to the ability to retain culturability on Columbia Blood Agar (CBA) containing 5% (v/v) defibrinated horse blood after exposure to sterile distilled water at two temperatures: 25°C and 4°C (Figure 1). At 4°C (72 h), whereas only approximately 17% of strain M1 cells were still culturable, for strains 1336 and 414 much higher levels (64% and 48% respectively) remained culturable. At 25°C (24 h), whereas only approximately 1.2% of strain M1 cells remained culturable, for strains 1336 and 414 approximately 71% and 82% respectively remained culturable. At these two sampling points, the survival of strain M1, based on culturability on CBA media, was significantly lower (p < 0.01; 2-tailed Student’s t-test) than for either of the other two strains.

**Figure 1.**
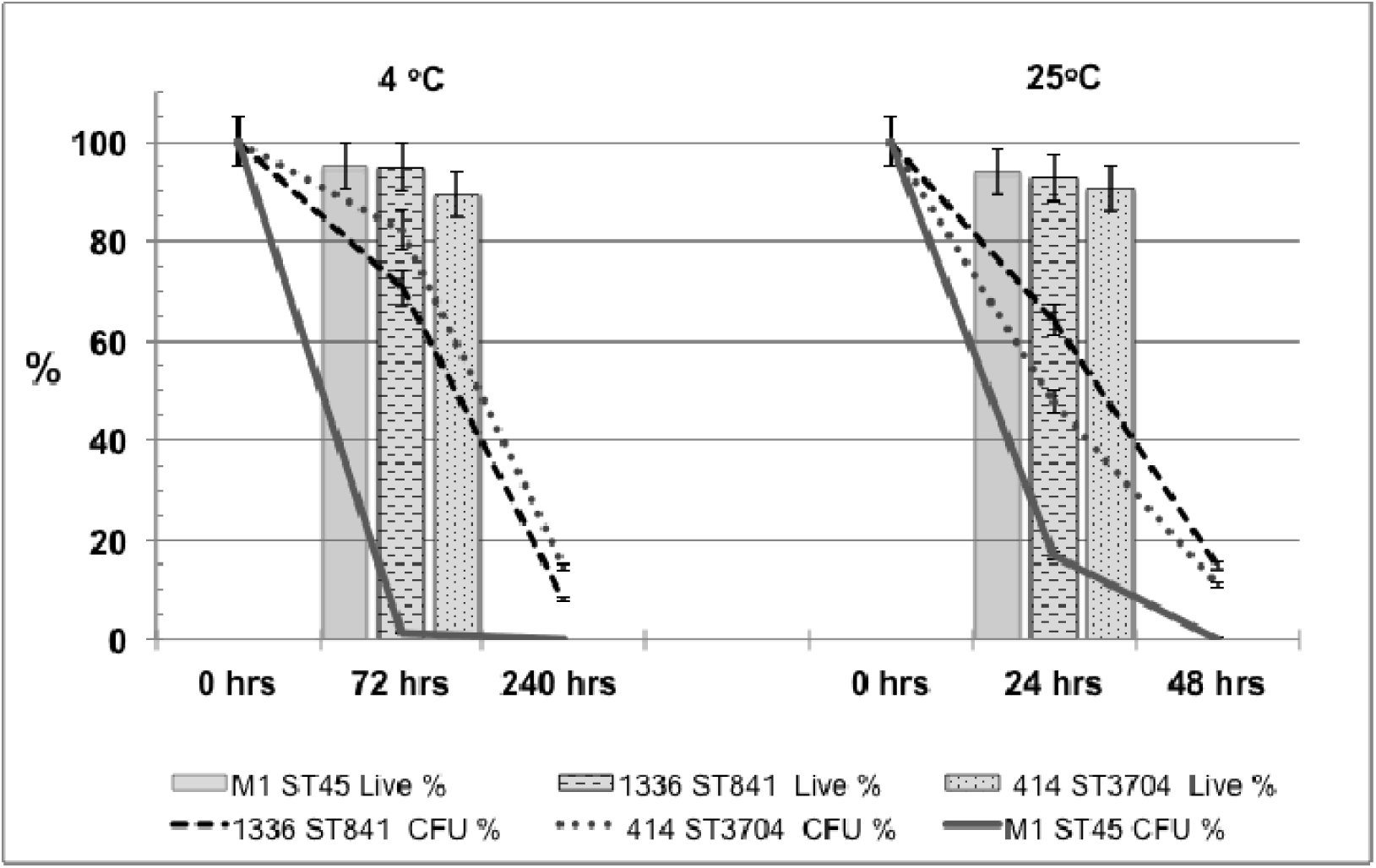
Culturability of *C. jejuni* strains during survival periods in sterile distilled water. The percentage of the original inoculum retaining culturability is shown at 0 h, 72 h and 240 h at 4°C and at 0 h, 24 h and 48 h at 25°C. The percentage of viable cells at 72 h at 4°C and 24 h at 25°C (indicated by the line graphs) was determined by LIVE/DEAD staining (BacLight, Invitrogen) for *C. jejuni.* The error bars represent a 5% error, three independent biological replicates were performed.

We further investigated viability of cells using LIVE/DEAD staining at Time 0, 4°C (72 h) and 25°C (24 h). Whereas the prevalence of culturable cells on CBA media declined rapidly, especially for strain M1, at both 4°C and 25°C, the percentage of viable cells according to LIVE/DEAD staining remained high throughout the experiment (Figure 1). Hence, although strain M1 rapidly loses culturability on CBA media during exposure to water, the majority of cells remain viable, indicative of the VBNC state when this medium is used.

### Overview of differential gene expression during survival of *C. jejuni* M1 in water

In order to better understand the process by which strain M1 remains viable but loses culturability, we analysed the transcriptome of cells recovered from water survival experiments. *C. jejuni* M1 gene expression in water was investigated, using Illumina RNA Seq under two key experimental conditions: i) 25°C for 24h, ii) 4°C for 72 h, selected because of the high viability but low culturability exhibited by strain M1 at these sampling points (Figure 1). In addition, two controls were included in the study. The first control involved suspending an inoculum of *C. jejuni* M1 in 100 ml of sterile distilled water and recovering the cells immediately; this control will be referred to as the “Time 0” sample. It should be noted that the bacterial cells in the Time 0 sample were pre-exposed to water at room temperature for approximately 20 min due to the sample processing time, prior to RNA extraction. The second control consisted of cells cultured in Mueller Hinton Broth (MHB) for 24 h at 37°C under microaerophilic conditions; this transcriptome will be referred to as the “MHB Control”. All initial inocula were taken from the same starter culture.

The transcriptomics data were analysed using two separate approaches to determine differential expression of genes: (i) based on counts per million (cpm) and (ii) using BitSeq analysis.

A summary of all M1 RNA Seq data in counts per million (cpm) is shown in Additional file 1. It is apparent from these data that significant transcriptome changes occurred during the preparation of the Time 0 sample compared to the MHB Control, indicating that *C. jejuni* gene expression changes in response to suspension in water occur rapidly. Figure 2 shows comparisons of genes up (Figure 2A) or down (Figure 2B) regulated 2-fold or more relative to the MHB Control during the different experimental conditions. Additional file 2 summarizes genes that are up- or downregulated 2-fold or more across all three test conditions compared to the MHB control.

**Figure 2.**
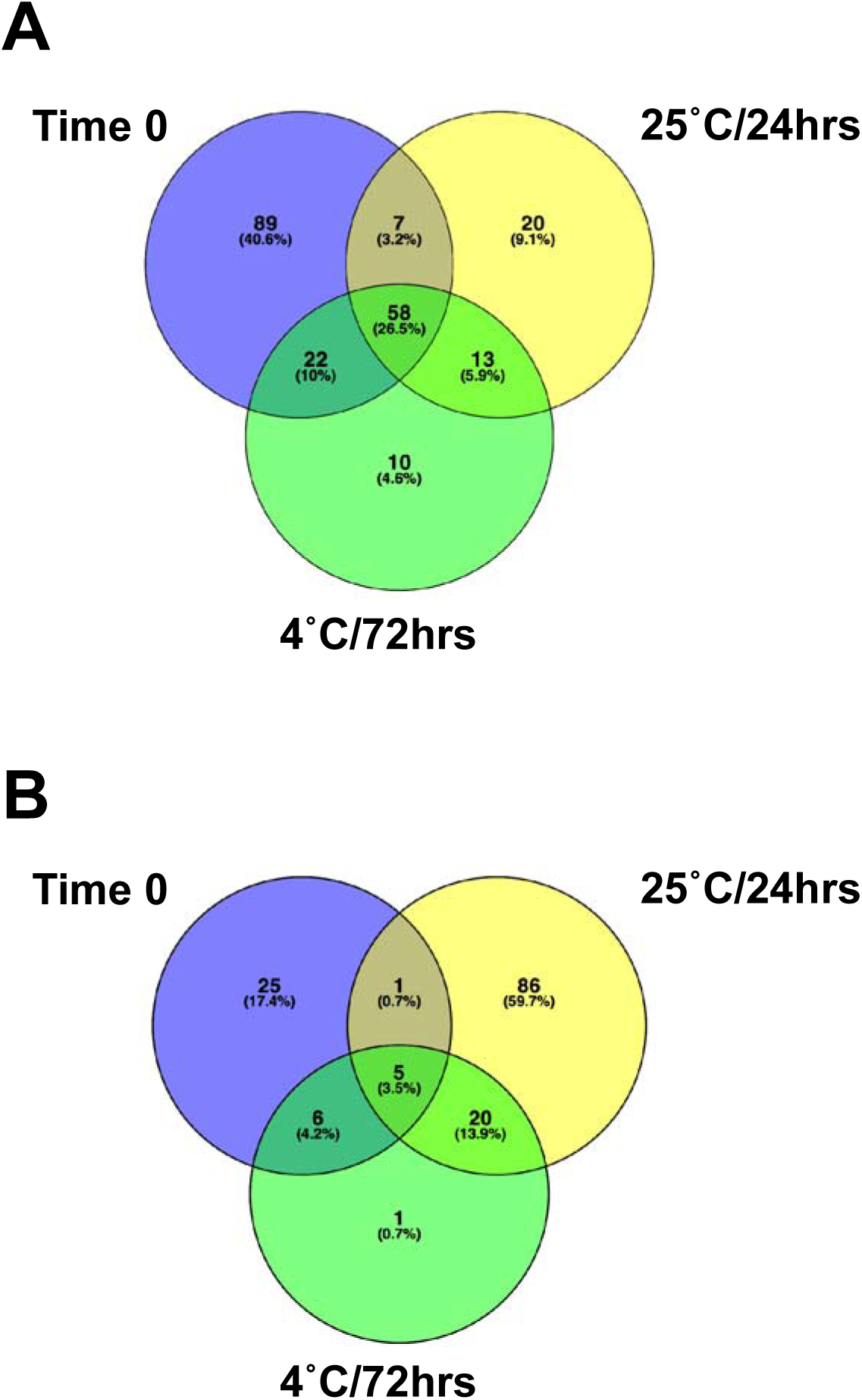
Summary of gene expression changes. (A) Venn diagram of Genes upregulated ≥ 2-fold (log counts per million) compared to the MHB control. (B) Venn diagram of Genes downregulated ≥ 2-fold (log counts per million) compared to the MHB control.

Figure 2 clearly shows that more genes were upregulated (58 genes) across the three time-point/temperature combinations [Time 0, 25°C (24 h) and 4°C (72 h)], than downregulated (5 genes). The biggest response in terms of upregulation in gene expression was observed during the early exposure to water at Time 0. The biggest response in terms of downregulation occurred at 25°C (24 h) (Figure 2).

Among genes upregulated across all three conditions were several flagellar genes (*flgBCDEGHF*J and *flaG*), as well as the *flmA* (*pseB*) gene, which is also involved in flagellar assembly and glycosylation; *flmA* mutants are non-motile and accumulate intracellular flagellin [40]. Four of the genes upregulated across all three conditions were hypothetical ones with their function currently unknown (Additional file 2). The list of genes downregulated 2-fold included three members of the *nap-*operon.

The results of the Bayesian Inference of Transcripts from Sequencing data (BitSeq) [41] analysis are summarized in Additional file 3. 546 genes were statistically significantly upregulated at Time 0, compared to the MHB Control, whereas 204 genes were downregulated; 872 genes showed no significant change. In the survival experiments, at 25°C after 24 h in sterile distilled water, 161 genes were upregulated, 557 genes were downregulated and 904 showed no significant change, compared to the MHB Control. At 4°C after 72 h, 202 genes were upregulated, 301 were downregulated and 1119 showed no significant change in expression compared to the MHB Control.

### Differential expression of known stress response genes

Despite its lack of many of the conventional pathways possessed by other enteropathogenic bacteria, *Campylobacter* does adapt quickly to environmental stressors [42, 43]. During adaptation to water, *C. jejuni* M1 must potentially counter oxidative and short term aerobic stress, hypo-osmotic stress, starvation and temperature shock in the different experimental conditions tested. Additional file 4 summarises the expression profiles of a number of previously characterized stress response genes compared to the MHB Control, across the experimental conditions tested. Figure 3 highlights the expression of known stress response genes and also includes all of the genes that showed significant up- or down-regulation (≥2-fold) compared to MHB Control (Additional file 2).

**Figure 3.**
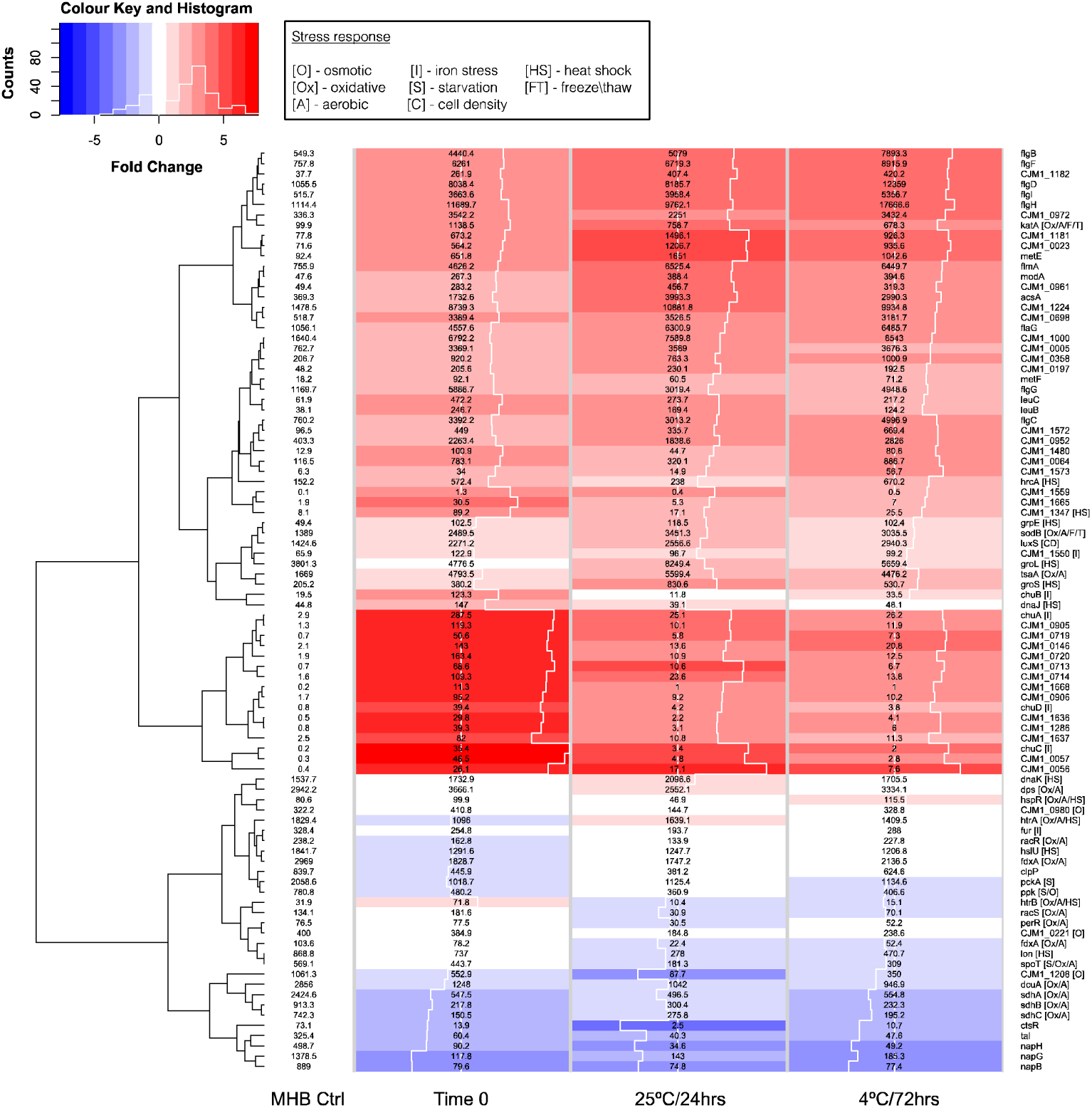
Expression changes in stress-related genes. The heatmap shows log fold change (logFC) calculated in edgeR compared to the MHB Control (blue = negative, red = positive, the white traceline in the columns indicates the size of the logFC measurement), and displays normalized counts per million (CPM) in numbers. It includes known stress response genes and genes up- or down-regulated >2-fold compared to the MHB control based on normalized CPM (numbers displayed in the heatmap matrix) for MHB Control, Time 0, 24 h (25°C) and 72 h(4°C). Square brackets display the type of stress response the gene is involved in (if known). The Histogram in the Colour key indicates the distribution of the data included in the heatmap.

### Oxidative, short term aerobic and temperature stress responses

Catalase *(katA*) and superoxide dismutase (*sodB*) genes were significantly upregulated in all three experimental conditions compared to the MHB control (Figure 3, Additional file 4). These genes are involved in both oxidative stress and freeze-thaw stress response [44]. Despite an apparent lack of cold shock proteins, *Campylobacter* maintains its metabolic rate at low temperatures (4°C) and appears to survive better at 4°C than at 25°C [45, 46]. In addition, an ankyrin-containing protein CJM1_1347 (Cj1386) has recently been identified in *C. jejuni,* encoded by a gene based directly downstream of the *katA* gene, and this is thought to be involved in the same detoxification pathway as catalase [47]; our data indicated significant upregulation of this gene as an early response to exposure to water in the Time 0 sample (Figure 3).

Previous studies have suggested that expression of the catalase (*katA*) gene, increased after exposure to oxidative stress, but not the superoxide dismutase (*sodB*) gene, the main antioxidant defence of most organisms [48]. It is essential for *C. jejuni* to counter atmospheric oxygen tensions and the resulting damage to nucleic acid through the toxicity of reactive oxygen species (ROS).

Unlike the three classified types of superoxide dismutases (SODs) present in *Escherichia coli, C. jejuni* only possess one (SodB) [49, 50]. It has been suggested that both SodB and catalase may play an important role in intracellular survival of *C. jejuni* [51, 52].

The *htrA* gene was significantly downregulated during the early response to exposure to water (Time 0). However, downregulation was not evident at the other time points. HtrA is important for stress tolerance and survival of Gramnegative bacteria generally [53, 54] and is required for both heat and oxygen tolerance [55]. It has been reported previously that *htrA* is downregulated in *C. jejuni* in response to low nutrient and especially oxidative stress [56]. HtrA, a periplasmic serine protease, displays both chaperone and protease properties, both implicated in the ability to tolerate stress, though the chaperone activity was identified as more important for resistance to oxidative stress [57].

*Campylobacter* cells show signs of heat stress at temperatures of 46°C and above; these conditions accelerate the transition of spiral cells to coccoid shaped ones. Arguably our experimental conditions tested for cold shock conditions, encountered by *Campylobacter* in the natural environment outside the host, with test temperatures below the ideal growth temperatures of 37 to 42°C. We did, however, observe changes in the expression levels of several heat shock proteins at 25°C including the chaperones *groLS* and *dnaK* the encoding genes are summarized in Additional file 4.

### Osmotic stress

While a number of studies have contributed to our understanding of hyperosmotic stress responses [58-60], mainly related to food preservation and *in vivo* environments, hypo-osmotic stress responses are still poorly understood. Introducing the bacteria into water (Time 0) leads to a marked influx of water into the cells due to the osmotic gradient. Hence, *C. jejuni* needs to react quickly to prevent cell lysis. Genes implicated in responding to hyper-osmotic conditions include *htrB, ppk* and a sensor histidine kinase (CJM1_1208). Both CJM1_1208 and *ppk* were downregulated at all three time points (Figure 3, Additional file 4).

The obvious response to changes in osmolarity, exhibited by many bacteria, is to pump out both water and solutes. Three stretch-activated mechano-sensitive (Msc) channels have been described in *E. coli:* MscM (mini), MscS (small) and MscL (large). A homologue for MscL has been described in *H. pylori* [61-63]. Kakuda *et al.* recently identified two putative mechanosensitive channels in the strain 81-176 (*Cjj0263* and *Cjj1025*), corresponding to CJM1_0221 (mechanosensitive ion channel family protein) and CJM1_0980 (putative membrane protein) respectively in M1 [64]. Both genes were upregulated during the early response at Time 0, compared to the MHB control, with CJM1_0980 showing statistically significant upregulation (PPRL=0.965). Interestingly, both genes were downregulated at 4°C (72 h), suggesting that *C. jejuni* M1 had adjusted to the hypo-osmotic conditions after 24 h (Figure 3).

### Iron acquisition

Iron acquisition is a vital process for bacterial survival and persistence. Due to the toxic potential of free iron, storage and uptake are tightly regulated. The putative hemin uptake gene cluster *chuABCD* and *Cj1613c* (*CJM1_1550*) are regulated by the ferric uptake repressor (Fur), which in turn is governed by the availability of free iron [65]. In this study, the *chuABCD* genes were upregulated >2 fold across all three experimental conditions tested, compared to the MHB control; *chuA* and *chuC* were statistically significantly upregulated over all three conditions, whereas *chuB, chuD* and the heme oxygenase gene *CJM1_1550* (*Cj1613c*) were only upregulated at Time 0. These findings strongly support the importance of iron regulation during water survival. *The fur* gene was statistically significantly downregulated at 25°C (24 h) compared to the MHB control, but did not vary significantly between the non-control samples.

### Starvation response

The stringent response is rapidly induced during starvation and stationary phase. Classically, RpoS is the global regulator for the stationary phase in bacteria. In lacking the *rpoS* gene, *C. jejuni* presents an RpoS-independent response to starvation and stationary phase. Inorganic polyphosphate (poly-P) is a linear polymer of phosphate residues linked by phosphoanhydride bonds which provide a high energy to the cell [66]. Poly-P, a source of energy and an essential molecule for survival during starvation, is synthesized by mediating the key enzyme polyphosphate kinase 1 (*ppk*) [67]; we observed that *ppk* was down-regulated in response to water (Figure 3; Additional File 4). Interestingly, it has been reported that a mutant *C. jejuni* (Δ*ppk*) showed decreased ability to enter a VBNC state due to lack in poly-P synthesis [16, 68]. Our observations of down-regulation appear to contradict this notion and may be indicative of variations between strains.

Even if glucose was available, *Campylobacter* spp. are incapable of using glucose as an energy source and have a very restricted carbohydrate catabolism (non-saccharolytic), a characteristic that distinguishes them greatly from other gastrointestinal pathogens. Phosphoenolpyruvate carboxykinase (PCK)(CJM1_0407), an essential enzyme in gluconeogenesis, was significantly upregulated in the early response (Time 0) compared to the MHB Control (Additional file 2).

### Quorum sensing

The transfer from an exponentially growing culture into water suddenly presents *C. jejuni* M1 cells with very low cell density conditions. Bacterial communities can communicate, in a density-dependent manner, through sensing autoinducers, extracellular signal molecules, produced by members of the community. The LuxS product autoinducer 2 (AI-2) is found in over 55 species, and is common to both Gram positive and Gram negative bacteria [69, 70]. Compared to the MHB control, we observed a significant upregulation of *luxS* expression at Time 0, and at both 25°C (24 h) and 4°C (72 h) (Additional file 2). The gene encoding CosR (CJM1_0334), a known positive regulator of *luxS* [71], was significantly upregulated at 4°C (72 h) compared to 25°C (24 h), but not found to be upregulated otherwise.

### Protein translocation and secretion

One of the overarching factors important in stress survival is the ability to transport proteins across membranes. The twin-arginine translocation (TAT) system in particular is vital for stress survival [72]. Our data show that components of the TAT system were upregulated at both Time 0 and at 4°C (72 h), compared to the MHB control (Figure 4), suggesting an increased need for translocation of proteins across the cell membrane at these time points and/or temperatures. In contrast, components of the Sec pathway were downregulated in all three conditions (Figure 4), confirming the crucial role of TAT during stressful conditions.

**Figure 4.**
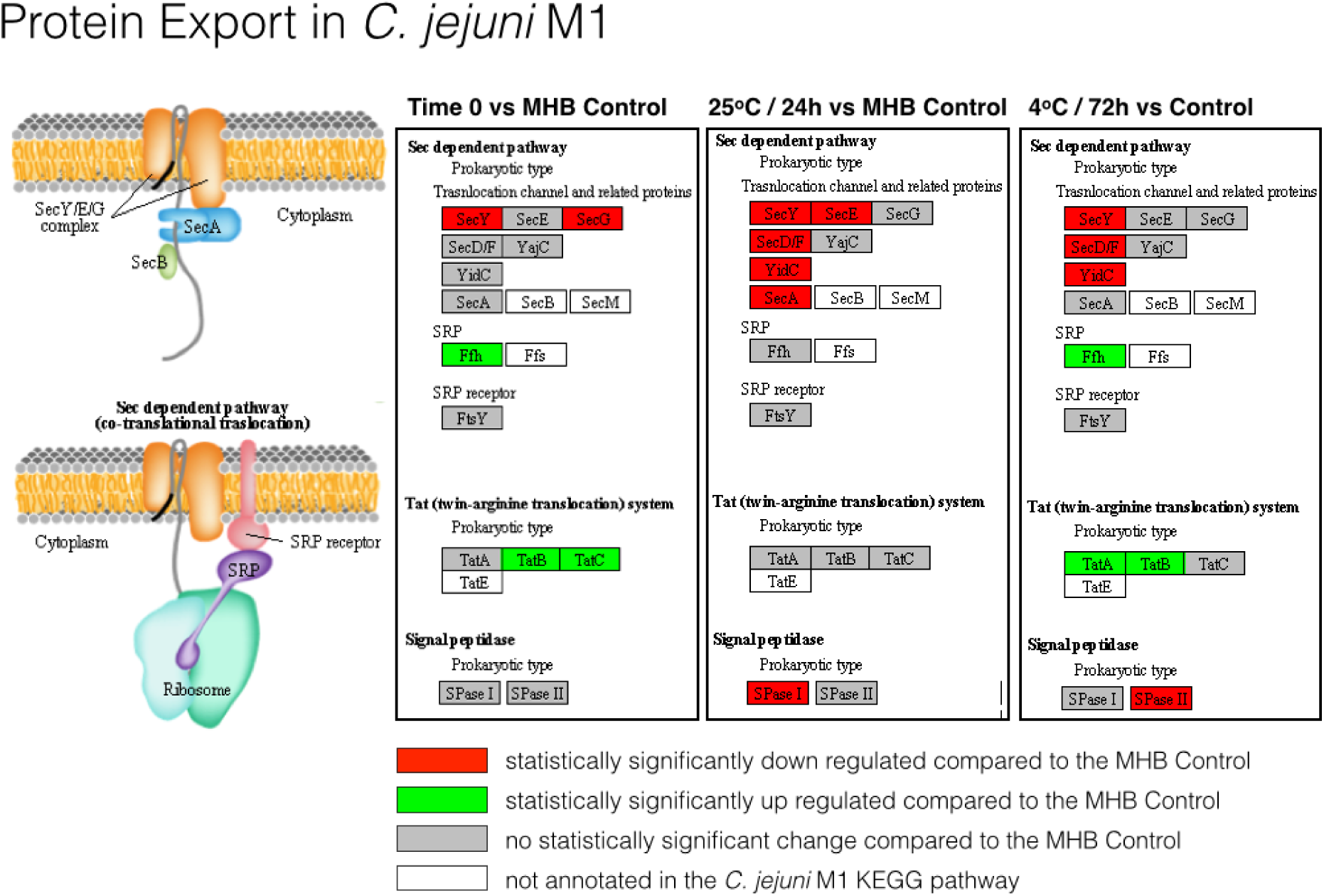
Significant expression changes determined through BitSeq analysis of Sec-dependent and twin-arginine (TAT) protein export pathway components mapped in KEGG. Results are shown for Time 0, 25°C (24 h) and 4°C (72 h) in sterile distilled water

The TAT system is an inner membrane translocase that transports proteins folded in the cytoplasm across the inner membrane. In contrast to Sec, which cannot accept tightly folded pre-proteins for translocation, the TAT system can translocate folded enzymes. Several substrates for the *C. jejuni* TAT system have been identified, including PhoX [73-76], the only alkaline phosphatase idenitified in *Campylobacter* species. Upon transport into the periplasm PhoX becomes active, providing *Campylobacter* with that vital energy source, inorganic phosphate (Pi). The gene encoding PhoX (CJM1_0145) was significantly upregulated at Time 0 compared to the MHB control.

### Motility and Chemotaxis

We observed the upregulation of many flagellar assembly genes across all experimental conditions (Figure 5); however, expression of both the flagellar motor genes and components of the chemotaxis pathway (Figure 6) was either downregulated or unchanged, indicating an alternative role to motility, as has been suggested previously [77]. The structural components of flagella are important for the secretion of virulence factors such as the *Campylobacter* invasion antigen (CiaB) [78]. However, under the conditions tested here, *ciaB* expression was not significantly upregulated (Additional file 3). It has been suggested that *C. jejuni* could also play an important role in adhesion in the early stages of biofilm formation [79]. Further work is needed to determine whether the gene expression changes that we have seen in water are indicative of the bacteria aggregating as a precursor to forming biofilms.

**Figure 5.**
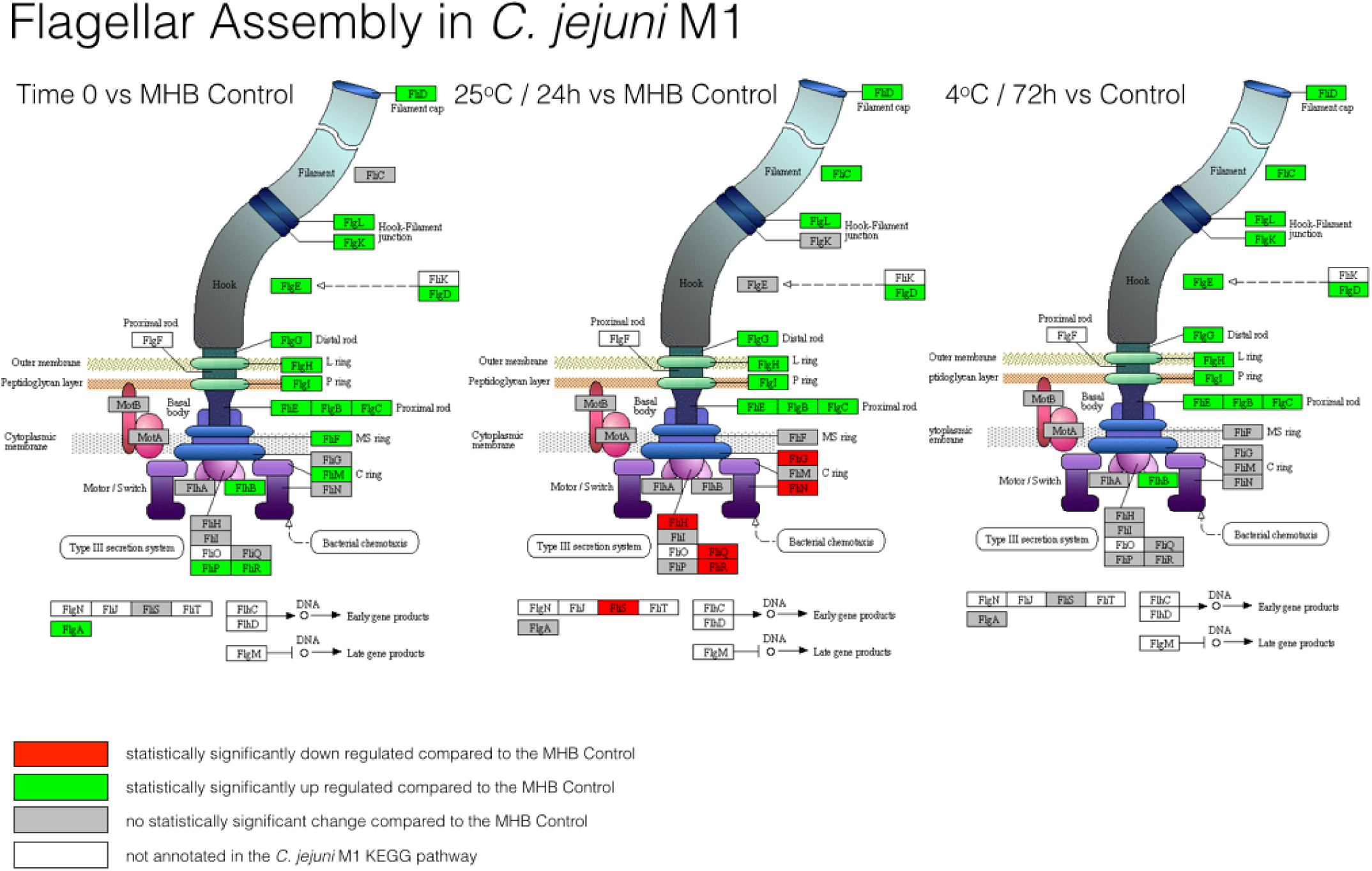
Significant expression changes determined through BitSeq analysis of *C. jejuni* M1 flagellar assembly components mapped in KEGG. Results are shown for Time 0, 25°C (24 h) and 4°C (72 h) in sterile distilled water.

**Figure 6.**
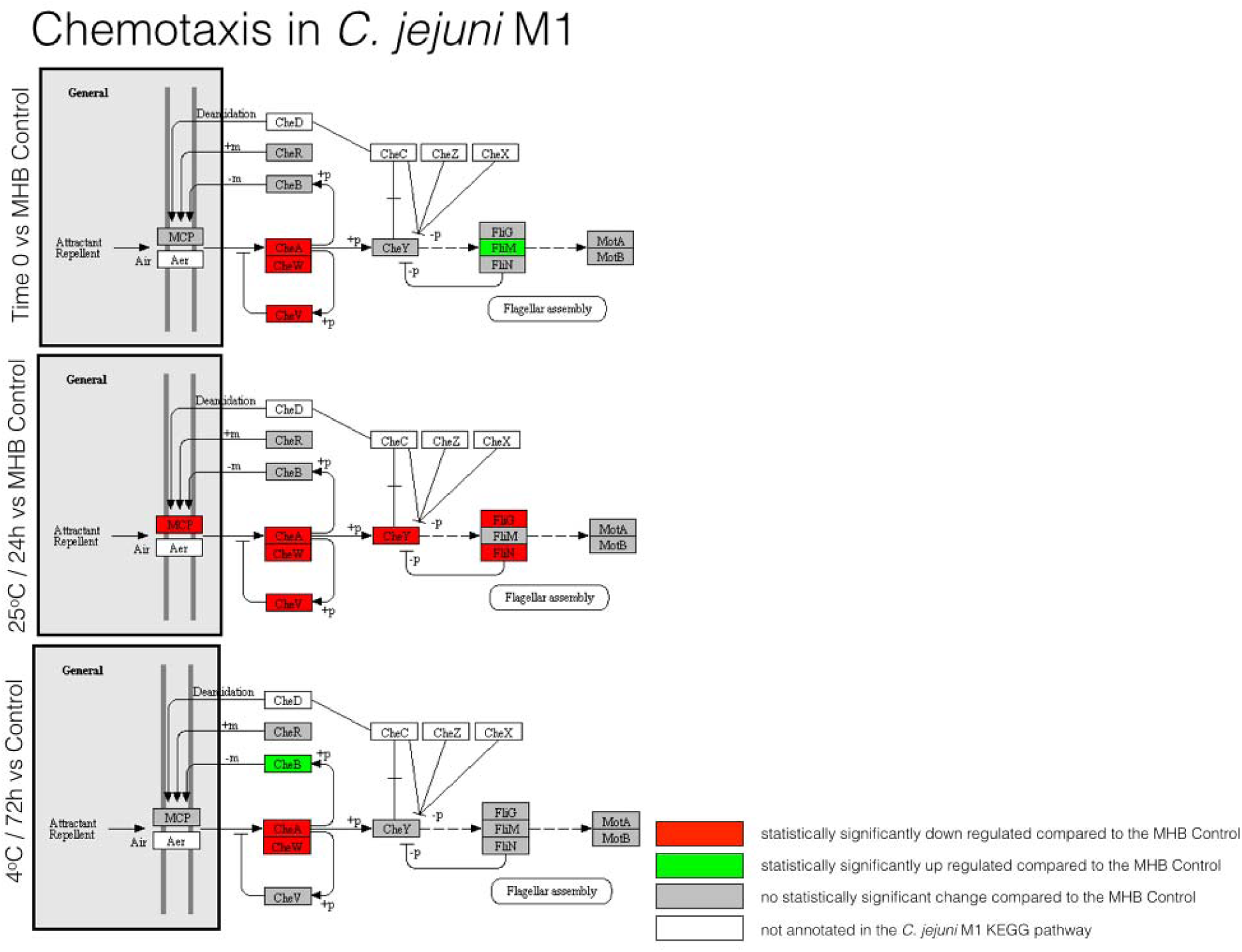
Significant expression changes determined through BitSeq analysis of *C. jejuni* M1 chemotaxis components mapped in KEGG. Results are shown for Time 0, 25°C (24 h) and 4°C (72 h) in sterile distilled water.

Studies in *H. pylori* have shown that the AI-2 encoded by *luxS,* which is highly upregulated across conditions in this study, targets expression of flagellar genes, and it has been suggested that *H. pylori* can regulate the composition of its flagella in response to environmental clues [80, 81]. Hence, there may be a link between the LuxS and flagellar component expression changes that we observed.

### Electron transport pathways and metabolism

The respiratory chain in *C. jejuni* is highly branched, with a number of potential electron acceptors, including fumarate, nitrate, nitrite, trimethylamine-N-oxide (TMAO) and dimethylsulphoxide (DMSO), involved in growth under severely oxygen-limited conditions [82]. In our study, three genes (*napG, napH, napB*) encoding enzymes that play an important role in major electron transport pathways were down-regulated in response to the water environment (Figure 3; Additional file 2). *nrfA* was also down-regulated in all three conditions, whilst *nrfH* was significantly down-regulated in two of the three conditions compared to the MHB control. These genes encode nitrate (Nap) and nitrite (Nrf) reductases involved in the use of nitrate and nitrite as electron acceptors. The Nap nitrate reductase is a two subunit enzyme comprising NapA and NapB, requiring NapD for proofreading. It is thought that the iron-sulphur proteins NapH and NapG assume the role of electron door to the NapAB complex [83]. The nitrite reductase NrfA is the terminal enzyme in the reduction of nitrite to ammonia, and is thought to play a role in defence against nitrosative stress [83]. It is thought that NrfH is the sole electron donor to NrfA [83]. Hence, our data suggest that nitrate or nitrite electron acceptors are not being utilised during survival in water.

The expression of many of the genes involved in central carbon metabolism was down-regulated in water. These included the genes encoding succinate dehydrogenase (SdhABC), malate dehydrogenase (Mdh) and formate dehydrogenase (FdhABC). Interestingly, the genes encoding PutP (proline transport) and PutA (proline dehydrogenase) were up-regulated, suggesting that proline metabolism was active. PutA catalyses the oxidation of proline to glutamate. Glutamine synthase (GlnA), which can catalyse conversion of glutamate to glutamine, and gamma-glutamyl transpeptidase (GGT), which can catalyse the hydrolysis of glutamine to form glutamate and ammonia, were also up-regulated. In *H. pylori*, it has been shown that the likely role of GGT is to supply the bacteria with glutamate for catabolism by the hydrolysis of extracellular glutathione or glutamine, with the substrates being hydrolysed in the periplasm before glutamate is transported into the cell [84]. Hence, there is some evidence for metabolism involving proline, glutamine and glutamate, but the full pathway is not clear.

### Pathogenicity and virulence factors

Unlike other enteropathogenic bacteria, *C. jejuni* does not possess many conventional pathogenicity factors. The cells in VBNC state may play an important role in the pathogenicity of *C. jejuni*; for example, the expression of the virulence gene *cadF,* encoding an outer-membrane protein of *C. jejuni* involved in adhesion to intestinal fibronectin has still been detected at high levels up to the third week of entering a VBNC state [85]. We observed consistently high levels of expression of *cadF* across all conditions tested (Additional file 1). However, we did not detect any statistically significant changes in *cadF* expression (Additional file 3). In contrast, the genes encoding cytolethal distending toxin (*cdtABC*) were down-regulated in response to water (Additional file 3).

### Do genomic Regions of Difference (RODs) in the *C. jejuni* M1 genome play a role in its enhanced survival in the experimental conditions tested?

To further investigate the water survival response of *C. jejuni* M1, we identified RODs within the *C. jejuni* M1 genome compared to the other *C. jejuni* strains tested in this experiment, *C. jejuni* 414 and 1336 (Figure 1 A and B), as well as the well characterized reference strain NCTC11168 [30, 38, 86]. We identified a total of 45 RODs in the *C. jejuni* M1 genome; 13 of these RODs are variable regions in all four genomes tested. The results have been summarized in Additional file 5 where shared RODs have been highlighted.

The currently published *C. jejuni* M1 genome (CP001900) contains 234 CDS annotated as “hypothetical protein” or “putative uncharacterized protein“. Albeit many of the functions and products of these genes have been described since, for ease of discussion we re-annotated the genome using a combination of PROKKA [87], and searches in stringDB and BLAST (BLASTP and BLASTX). Putative functions and expression profiles are summarised in Additional file 5. An overall summary of the M1 genome, including RODs, hypothetical genes, and BitSeq expression data is shown in Additional file 6.

There were a number of examples of genes, or clusters of genes, within RODs that were either up- or down-regulated (Additional file 5; Additional file 7). It has been reported previously that the genomes of strains 1336 and 414 lack some genes that are widely distributed in *C. jejuni* [30]. Hence, some RODs include genes that have known putative functions, such as the *cdtABC* genes, which are down-regulated in water. Since M1 behaves differently to strains 1336 and 414 in water, it is possible that genes present in only strain M1 might contribute to these differences. For example, ROD-6 (comprising four putative genes) is absent from strains 414 and 1336, and is upregulated in water. However, the gene functions are not known. Further work would be needed to determine which, if any, RODs play a significant role in the strain M1 VBNC phenotype.

## Conclusions

Our data suggest that *C. jejuni* M1 adapts rapidly to introduction into a water environment, instigating gene expression changes that allow it to adapt to the stressful conditions, whilst maintaining viability. In addition to the up-regulation of stress responses, a preference for secretion via the Tat pathway and the down-regulation of many (but not all) metabolic genes, the bacteria adapt in some more surprising ways. We observed down-regulation of chemotaxis genes coupled to up-regulation of flagellar genes, suggesting a role for flagella that is not linked to motility. In addition to the secretion of virulence factors, flagella have been implicated in autoagglutination and microcolony formation as a precursor to biofilm formation [88]. Hence, it is possible that the observed gene expression changes are indicative of the bacteria starting to instigate a lifestyle change from sessile to biofilm as a survival strategy. We also found evidence that the putative quorum sensing system protein LuxS plays a role during the adaptation to water. Genes in the accessory genomes, including genes encoding hypothetical proteins of no known function, may play a role in the variable survival phenotypes that are observed between strains. Hence, despite the loss of culturability, strain M1 remains viable and adapts to suspension in water via multiple specific changes in gene expression. Further work is needed to ascertain which responses are shared by all *C. jejuni* strains, and which are specific to a sub-set of strains sharing the characteristics of M1.

## Methods

### Bacterial Growth conditions

*Campylobacter* strains were stored on storage beads in glycerol broth (M Lab) at -80°C. *C. jejuni* strains were grown on sterile Columbia Blood Agar Base (CBA, Oxoid) with 5% (v/v) defibrinated horse blood (Oxoid) at 37°C for 48 h under microaerobic conditions (85% [v/v] N_2_, 5% [v/v] O_2_, and 10% [v/v] CO_2_), in a Whitley VA500 Workstation incubator (Don Whitley Scientific Ltd).

### Preparation of cell suspensions for testing survival in sterile water

For survival experiments, bacteria were sub-cultured on blood agar for 24 h at 37°C under microaerobic conditions. A 5 μL loop of cultured cells was taken, and suspended in 5 mL of Muller-Hinton Broth (MHB, Oxoid) supplemented with *Campylobacter* growth supplement (LAB M). Suspended bacterial samples were adjusted to a final optical density at 600 nm (OD_600_) of 0.05 [3.8×l0^7^ - 3.5×l0^8^ Colony Forming Unit (CFU)/mL] (Spectronic Biomate 5).

### Preparation and inoculation of sterile water sample

Filtered-tap water (PUR1TE SELECT) was collected, and autoclaved at 121°C for 15 min. Aliquots of 99 mL (pH 6.5) of autoclaved water samples were transferred into 250 mL sterile borosilicate glass bottles with screw caps (Schott, Duran, Germany) in triplicate. These were inoculated with 1 mL from bacterial suspensions to a final concentration of 8×l0^5^ - 3.7×l0^6^ cells/mL. The inoculated samples were kept in the dark at 25°C (for 24h) or 4°C (for 72 h). An uninoculated sterile distilled water sample for each temperature was used as a control for the presence of contamination. A further control was used whereby the water was inoculated with 1 mL of bacterial suspension and the bacteria were collected immediately (time 0). All experiments were conducted using three independent technical replicates and three biological replicates (strains M1, 1336 and 414).

### Enumeration of colony forming units (CFU)

At time 0, 25°C (24 h), and 4°C (72 h), a 100 μL sample was taken and 10-fold dilutions were made in MHB supplemented with *Campylobacter* growth supplement. A 10 μL spot assay of appropriate dilutions was carried out on CBA plates in triplicate. The plates were incubated for 48h at 37°C under microaerophilic conditions, and the survival was then determined by enumerating the CFU/ mL.

### Cell survival by LIVE/DEAD staining

Inoculated water samples were prepared as described above. At time 0, 25°C (24 h), and 4°C (72 h), inoculated water samples *C. jejuni* strains M1,414 and 1336 were concentrated by centrifugation at 3893 × g for 20 min (3-16PK - SIGMA) in Falcon tubes (Corning, Appleton Woods). Supernatant was removed and approximately 1 mL remained at the bottom of each Falcon tube and was subsequently transferred into a 1.5 mL Eppendorf tube. Cells were pelleted by centrifugation at 5000 × g for 20 min. The pellet was re-suspended in 1 mL of sterile distilled water. Using the LIVE/DEAD BacLight, Invitrogen kit, 3 μL of the mixture (SYTO 9 green-fluorescent and propidium iodide red-fluorescent) were added for each 1 mL of bacterial suspension and this mixture was incubated for 15 min at room temperature. 5 μL of the cell suspension was then placed on a microscope slide, covered with a 22 × 22 mm cover slip and sealed.

The LIVE/DEAD BacLight kit (Invitrogen) contains two nucleic acid stains, SYTO 9 dye, which penetrates live cells (intact membranes) causing the cells to stain fluorescent green, and propidium iodide dye that cannot cross the cell membrane and therefore only stains cells red if the membranes are damaged and the cell is therefore presumed dead.

Enumeration of VBNC cells was carried out under a fluorescence microscope (Nikon ECLIPSE 80i). For each sample, three fields were enumerated at an average of 90-180 cells in each field. The percentage of viable cells was calculated as follows: % viable cells = [viable cell count (green cells)/total cell count (green cells + red cells)] × 100. The experiment was conducted in three independent replicates.

### RNA extraction from water survival experiments

100 mL of inoculated water samples were prepared for *C. jejuni* M1 in triplicate at time 0, 25°C (24 h) and 4°C (72 h). Cells were concentrated by centrifugation at 3893 × g (3-16 pk-SIGMA) for 20 min at the corresponding temperature in 50ml Falcon tubes (Corning, Appleton Woods). Supernatants were removed but, for each sample, approximately 1 ml was retained at the bottom of each Falcon tube and subsequently combined per sample and transferred into a 1.5 mL Eppendorf tube. Cells were pelleted by centrifugation at 5000 × g for 10 min. A negative control was included using only sterile distilled water. Additionally a control of *C. jejuni* M1 grown in Mueller Hinton Broth was prepared. Cells were grown to a density of 1x10^7^ in a 25 cm^2^ cell culture flask with gas exchange lid (Corning) at 37°C microaerophilic conditions. Cells were collected by centrifugation at 3000xg in a microcentrifuge (Eppendorf). Once collected, cells were immediately resuspended in TRIzol solution (3 times TRIzol volume to 1 volume of cells) (Ambion) and stored at -80oC until further processing.

TRIzol samples were allowed to reach room temperature and cells were disrupted using vigorous vortexing. The samples were then incubated at room temperature for 5 min. RNA was extracted using the Direct-zol RNA MiniPrep Kit (Zymo Research), following the manufacturer’s instructions.

### Illumina Library construction and sequencing

116 ng of total RNA was depleted using the Illumina Ribo-zero rRNA Removal Kit (Bacteria) and purified with Ampure XP beads. Successful depletion was confirmed using Qubit and Agilent 2100 Bioanalyzer. All of the depleted RNA was used as input material for the ScriptSeq v2 RNA-Seq Library Preparation protocol. Following 15 cycles of amplification the libraries were purified using Ampure XP beads. Each library was quantified using Qubit and the size distribution assessed using the Agilent 2100 Bioanalyzer.

The final libraries were pooled in equimolar amounts using the Qubit and Bioanalyzer data. The quantity and quality of each pool was assessed by Bioanalyzer and subsequently by qPCR using the Illumina Library Quantification Kit from Kapa on a Roche Light Cycler LC480II according to manufacturer’s instructions. Sequencing was performed at the Centre for Genomic Research, University of Liverpool, on one lane of the Illumina HiSeq 2500 2x125 bp using v4 chemistry (Illumina).

### RNA Seq data analysis

The raw FASTQ data files were trimmed for the presence of Illumina adapter sequences using Cutadapt (v1.2.1.) [89], using the −0 3 option. The reads were further trimmed using Sickle (vl.200) [https://github.com/naioshi/sickle] with a minimum window quality score of 20. Reads shorter than 10bp after trimming were removed.

Sense and antisense overlaps between the annotation and mapped reads were counted using the HTSEQ package [90], using the stranded and union options. Read counts were then normalised and Differential Expression calculated using EdgeR implemented in R (version 3.1.2 (2014-10-31), using Loess-style weighting to estimate the trended dispersion values, heatmaps were constructed using heatmap2 in R.

For pairwise Differential Expression analysis between samples, the data were re-mapped to the *C. jejuni* M1 genome [38] using Bowtie2 [91], and parsed using the BitSeq (Bayesian Inference of Transcripts from Sequencing data) pipeline [41]. BitSeq takes into account biological replicates and technical noise, and thereby calculates a posterior distribution of differential expression between samples.

Statistical Significance of BitSeq results was visualized in Artemis [92]. Regions of Difference between the genomes of M1 (CP001900) [38], 1336 (ADGL00000000), 414 (ADGM00000000) [30] and NCTC11168 (AL111168) [86, 93] were derived through pairwise genome comparisons in ACT [94]. Putative / Hypothetical genes were selected in Artemis and a putative function was derived from searches using BLASTX (http://blast.ncbi.nlm.nih.gov/blast/Blast.cgi?PROGRAM=blastx&PAGE TYPE=Bl astSearch&LINK LOC=blasthome).

## Ethics Approval and Consent to Participate

Not applicable

## Consent for Publication

Not applicable

## Availability of data and materials

Sequence data are available at the European Nucleotide Archive under Primary Accession PRJEB17925. Other data supporting the results are available as Additional Files. Bacterial strains will be made available on request.

## Competing Interests

The authors declare that they have no competing interests.

## Funding

We acknowledge the Medical Research Council, Natural Environment Research Council, Economic and Social Research Council, Biotechnology and Biosciences Research Council and Food Standards Agency for the funding received for this project through the Environmental & Social Ecology of Human Infectious Diseases Initiative (Enigma; Grant Reference: G1100799/1).

## Author’ Contributions

CB carried out the transcriptomics experiments. KM, CEJ carried out the initial water survival experiments. CN and AL carried out the laboratory work required for library preparation and Illumina sequencing. CB and IBG analysed the data. TH, NW and CW conceived the study. CB and CW wrote the paper. All authors read and approved the final manuscript.

## Acknowledgements

We would like to thank John Kenny at the CGR for help and advice.

We thank members of the ENIGMA Consortium for helpful discussion. The ENIGMA Consortium investigators comprise Sarah O’Brien (PI), Rob Christley, Christiane Hertz-Fowler, Paul Wigley, Nicola Williams and Craig Winstanley (University of Liverpool), Peter Diggle (Lancaster University), Iain Lake, Kevin Hiscock and Paul Hunter (University of East Anglia), Ken Forbes and Norval Strachan (University of Aberdeen), Rachel Griffith and Dan Rigby (University of Manchester), Paul Cross (Bangor University), Stephen Rushton (Newcastle University), Tom Humphrey (Swansea University), Malcolm Bennett (University of Nottingham), David Howard (Centre for Ecology and Hydrology), and Brendan Wren (London School of Hygiene and Tropical Medicine).

## Additional files

**Additional file 1.** Summary of normalised expression values for each gene in average counts per million (cpm) determined in edgR.

**Additional file 2.** Table summarizing genes that are up or downregulated ≥ 2fold compared to the MHB control in read counts per million (cpm) across all three experimental conditions. Cpm displayed are normalised by the Trimmed Mean of M-values (TMM) method, implemented in the edgR Bioconductor package. In brackets the log-fold-change cpm relative to the MHB is shown. Cells are shaded according to log-fold-change (yellow=negative, green=positive).

**Additional file 3.** Summary of Bayesian Inference of Transcripts from Sequencing data (BitSeq) analysis [41] results. Results are shown in Probability of Positive Log Ratio (PPLR), PPLR ≥0.95 shows a high probability that the transcript is upregulated (green shading) compared to the first condition; PPLR ≤0.05 shows a very low probability that the transcript is upregulated compared to the first condition and therefore the probability of the transcript being downregulated (red shading) is very high.

**Additional file 4.** Table showing gene expression of previously described stress response genes shown as average read counts per million (cpm). Fold change compared to the MHB Control, as determined by edgeR, the Paired student’s T-test and PPLR determined through BitSeq analysis are shown in brackets. Stressor abbreviations: S, starvation; O, osmotic; OX, oxidative; A, aerobic; HS, heat-shock; F/T; freeze-thaw; I, iron stress; CD, cell density

**Additional file** 5. Regions of difference (RODs) identified between *C. jejuni* M1, 414,1336 and NCTC11168. RODs shaded in teal are divergent from M1 in the other three strains (414, 1336 and NCTC11168)

**Additional file 6. The *C. jejuni* M1 genome and maps a global overview of gene expression changes under the different test conditions in *C. jejuni* M1** From the outside in: Track 1 *C. jejuni* M1 genome; Track 2 CDS forward strand; Track 3 CDS reverse strand; Track 4 hypothetical/ putative uncharacterized genes (CDS); Track 5 and 6 Regions of Difference (RODs) in the *C. jejuni* M1 genome compared to *C. jejuni* 414 (turquoise), M1 compared to NCTC11168 (magenta) and M1 compared to 1336 (blue); Track 7 operons in M1 as predicted by Rockhopper; Track 8 genes that are statistically significantly upregulated (green) or downregulated (red) at Time 0 only, compared to the Control; Track 9 genes that are statistically significantly upregulated (green) or downregulated (red) in 25°C (24 h) only compared to the Control; Track 10 genes that are statistically significantly upregulated (green) or downregulated (red) in 4°C (72 h) only compared to the Control; Track 11 genes that are statistically significantly upregulated (green) or downregulated (red) in 25°C (24 h) only compared to Time 0; Track 12 genes that are statistically significantly upregulated (green) or downregulated (red) in 4°C (72 h) only compared to Time 0; Track 13 genes that are statistically significantly upregulated (green) or downregulated (red) in 4°C (72 h) only compared to 25°C (24 h). Colouring of Tracks 2 and 3: replication related = bright red; efflux pumps = bright green; Chemotaxis = bright blue; hydrogenases = pale pink; iron or heme-related = light brown; hydrogenases = rose; chaperone = orange; lipoproteins = yellow; membrane or periplasmic proteins = turquoise; ATP-/ ABC transporters = light green; ribosomal/RNA/ribonuclease = light blue; flagellar-related = magenta; hypotheticals = salmon

**Additional file 7.** Summary of hypothetical genes in the *C. jejuni* M1 genome and the corresponding BitSeq data. Green shading indicates PPLR ≥0.95, red shading indicates ≤0.05, grey shading indicates no statistically significant difference.

**Additional file 8.** Gel electrophoresis to show the quality of RNA samples. Example samples are shown for a sample from 25 °C (24 h) (D1 3), Time 0 (D0 2), 4 °C (72 h) (D3 1) and the Mueller Hinton Broth Control (C1 107). There was no evidence for RNA degradation in the 24 h or 72 h water samples compared to the time zero or control samples.

